# Vowel-like patterns modulate auditory P100m response but not its association with language abilities in children with ASD

**DOI:** 10.1101/2025.04.10.648101

**Authors:** Fadeev Kirill.A., Romero Reyes Ilacai V., Goiaeva Dzerassa E., Obukhova Tatiana S., Ovsiannikova Tatiana M., Prokofyev Andrey O., Rytikova Anna M., Novikov Artem Y., Stroganova Tatiana A., Orekhova Elena V.

## Abstract

**Background:** The P100/P100m component of auditory event-related potentials/fields is considered a potential biomarker of atypical arousal and language abnormalities in children with ASD. When elicited by complex speech-like sounds with regular temporal or frequency structure, P100/P100m may be influenced by sustained negativity (SN), which can reduce its amplitude due to opposing current polarity.

**Methods:** Using MEG, we investigated how acoustic regularities affect P100m latency and amplitude differences between TD children and those with ASD. MEG was recorded in 35 ASD and 39 TD boys (7–12 years) in response to control sounds (non-periodic, non-vowels) and stimuli with temporal regularity (periodic non-vowels), frequency regularity (non-periodic vowels), or both (periodic vowels). P100m was estimated using distributed source localization.

**Results:** In both groups, P100m amplitude and latency decreased in response to acoustic regularities, accompanied by a proportional increase in SN. No group differences were observed in P100m latency, amplitude, or their modulation by stimulus characteristics. In ASD, P100m latency variability was increased, and higher P100m amplitudes in the left auditory cortex were negatively associated with cumulative language and intellectual abilities.

**Conclusions:** In children, changes in P100m in response to acoustic regularities are most parsimoniously explained by an enhancement of SN with opposite polarity. No consistent relationship was found between P100m parameters or their modulation by acoustic regularities and ASD diagnosis. However, variations in cortical maturation and/or habituation processes, which affect the left-hemispheric P100m, may be relevant to cognitive and language functioning in children with ASD.

## 1. Introduction

Advances in understanding the neurobiology of neurodevelopmental disorders, coupled with progress in developing corresponding animal models, have highlighted the need for accessible neural biomarkers that are translatable between humans and animals. In this context, transient components of auditory cortical evoked response are of particular interest, as they can be reliably recorded in both animal models and human populations—including those with severe impairments, such as children with fragile X syndrome (FRAX) (An et al., 2022) or Rett syndrome (Kostanian et al., 2023; see Sysoeva et al., 2020 for review).

Maturation of the auditory cortex is associated with marked changes in the morphology of transient auditory evoked responses recorded by MEG and EEG. Unlike adults, who exhibit a P50-N100-P200 waveform in auditory ERP at vertex recording sites, young children typically display a P100-N200 complex (Ponton et al., 2000; Ponton et al., 2002). Both the latency and amplitude of the P100 component decrease with age between approximately 6 and 12+ years (Sharma et al., 1997; Ponton et al., 2000), likely reflecting the developmental emergence of the N100 component associated with maturation of supragranular cortical layers II and III (Eggermont and Moore, 2011). Some authors interpret this vertex-positive pediatric component with peak latency of ∼100 ms as an immature P50 (or M50 in MEG). However, since in MEG recordings, the earlier P50/M50 component of lower amplitude can be distinguished from a later (∼100 ms) component of the same polarity under some stimulation conditions (Orekhova et al., 2013), we adopt the P100/P100m designation here.

P100/P100m is a pre-attentive component that can be reliably detected in most children. Its developmental changes reflect the maturation of the auditory cortex (Eggermont & Ponton, 2003; Eggermont & Moore, 2011), making it a useful marker for assessing auditory cortex plasticity in cochlear implant users implanted at various ages (Sharma et al., 2015). Multiple studies have also evaluated characteristics of this component in children with developmental disorders characterized by atypical auditory sensory processing, such as autism spectrum disorders (ASD) (Cotter et al., 2023; Donkers et al., 2015; Haesen et al., 2011), Rett syndrome (Kostanian et al., 2023; Brima et al., 2024) or FRAX (An et al., 2022). Some research has identified relationships between P100m characteristics and children’s language abilities (Yoshimura et al., 2021; Matsuzaki et al., 2020; Kautto et al., 2024; An et al., 2022; Roberts et al., 2019), suggesting that P100m (M50) latency could serve as a biomarker for language abnormalities in ASD and possibly other developmental disorders (Roberts et al., 2019, 2021). Thus, P100m may reflect not only structural maturation of the auditory cortex but also its functional efficiency in speech processing.

The P100m and other transient components of the auditory response may overlap with the vertex-negative sustained response, referred to as SN (sustained negativity), which is observed throughout the entire duration of the auditory stimulus presentation when using low high-pass filter values or DC recording (Picton et al., 1978). Despite developmental differences in transient components, SN can be recorded in both children and adults (Stroganova, 2020; Orekhova et al., 2024). The magnitude of the SN depends on the properties of the auditory stimulation, being stronger for auditory patterns such as periodic stimuli (Keceli et al., 2012, 2015; Gutschalk et al., 2002, 2004; Gutschalk & Uppenkamp, 2011; Orekhova et al., 2024), or ‘sound figures’ characterized by either the presence of vowel formant structure (Hewson-Stoate et al., 2006; Orekhova et al., 2024; Gutschalk & Uppenkamp, 2011) or non-vocal frequency regularities (Molloy et al., 2019).

The brain rapidly detects the presence of temporal or frequency regularities, leading to an increase in SN as early as ∼40 ms post-stimulus (Molloy et al., 2019; Gutschalk et al., 2004). This early onset may reflect the activation of auditory cortical neurons involved in detecting potentially relevant sound patterns (Walker et al., 2011). Sustained negativity, rapidly increasing after its onset, can affect the properties of the P100/P100m component depending on the type and degree of regularity of the auditory pattern. However, the influence of auditory patterns on transient components of the auditory response in children has not been systematically investigated. If such an influence exists, intersubject differences in P100/P100m - including those associated with ASD - could potentially be affected by the auditory cortex’s capacity to automatically detect such patterns. Given the language deficits common in children with ASD and other developmental conditions, it would be important to explore how speech-inherent acoustic patterns, such as periodicity (pitch) or formant structure, modulate the P100 component of the auditory evoked potentials.

Previous MEG studies in children showed that, compared to the non-periodic non-speech complex sounds, sounds characterized by F0 periodicity, formant composition, and especially their combination in computer-generated vowels elicit an early (<100 ms) increase in SN in auditory cortical areas - the ‘sustained processing negativity’ (SPN) (Orekhova et al., 2024; Fadeev et al., 2024) - which may potentially affect children’s P100m. *The first aim* of the present study was to investigate how vowel-characteristic regularities influence the P100m component in children’s auditory cortex. Our *second aim* was to examine whether P100m parameter differences between children with ASD and typically developing (TD) children are influenced by these regularities. Since previous research has associated P100m latency (Yoshimura et al., 2021; Matsuzaki et al., 2020; Roberts et al., 2019) and amplitude (Yoshimura et al., 2013; Yoshimura et al., 2024; Kautto 2024; Roberts et al., 2019) with language abilities in children, our *third aim* was to explore relationship between these P100m characteristics and overall language skills in children with ASD.

## 2. Method

### 2.1 Participants

The study included 39 boys with ASD aged 6.9 – 13.0 years and 35 typically developing (TD) boys matched for age range. These participants were the same cohort as in the study investigating the sustained auditory MEG response (Fadeev et al., 2024). TD children were recruited via media advertisements. None of the TD participants had any known neurological or psychiatric disorders. Participants with ASD were recruited through multiple sources, including media advertisements, consulting centers, and an educational center affiliated with the Moscow State University of Psychology and Education.

The ASD diagnosis was based on the Diagnostic and Statistical Manual of Mental Disorder (5th ed.) criteria and confirmed by a child psychiatrist (author N.A.Y.) through clinical interviews with the child and parents/caregivers. Intellectual abilities were assessed using the KABC-II test, with the Mental Processing Index (MPI) serving as an IQ equivalent (Kaufman & Kaufman, 2004). The MPI excludes tasks requiring verbal reasoning, verbal concepts, or cultural knowledge, making it suitable for evaluating nonverbal abilities in both TD children and those with ASD (Drozdick et al., 2018).

The hearing status of all participants was assessed using pure–tone air conduction audiometry with an AA-02 clinical audiometer (“Biomedilen”). Auditory thresholds were tested at 500, 1000, 2000, 4000 Hz, and the average threshold cross these four frequencies was calculated separately for each ear. All participants demonstrated normal hearing (threshold< 20 dB HL) in both ears (British Society of Audiology, 2018).

The Ethics Committee of the Moscow State University of Psychology and Education approved this study. All children provided verbal assent to participate in the research, and their caregivers provided written informed consent for participation.

### 2.1 Assessment of the general level of language development

Language abilities were assessed using the Russian Child Language Assessment Battery (RuCLAB; Ivanova et al., 2016), measuring expressive and receptive skills in vocabulary (word production/comprehension) and morphosyntax (sentence production/comprehension). Test administration and scoring procedures followed that described in Fadeev et al. (2024). Discourse production subtests were excluded from this analysis, as most children with ASD could not complete them. To quantify overall language proficiency, we calculated the arithmetic mean of all remaining subtest scores, hereafter termed the *total language score*.

### 2.2 Stimuli

The experimental paradigm used in the present study is identical to that described in Fadeev et al. (2024). We used four types of synthetic vowel-like stimuli previously employed by Uppenkamp et al. (2006) and downloaded from ‘‘http://medi.uni-oldenburg.de/members/stefan/phonology_1/”. Five strong vowels were used: /a/ (caw, dawn), /e/ (ate, bait), /i/ (beat, peel), /o/ (coat, wrote) and /u/ (boot, pool).

The synthetic periodic vowels consisted of damped sinusoids, which were repeated with a 12 ms period, resulting in fundamental frequency of 83.3 Hz for each vowel. The carrier frequencies of each vowel remained fixed at the four lower formant frequencies selected within the typical range of an adult male speaker. These stimuli are hereafter referred to as periodic vowels. These regular vowel stimuli were modified, as described below, to create three additional stimulus classes: non-periodic vowels, periodic non-vowels, and non-periodic non-vowels. To disrupt periodicity, the start time of each damped sinusoid was jittered within ±6 ms relative to its start time in the original vowel, separately for each formant. Despite resulting in degraded voice quality (hoarse voice), these non-periodic sounds were still perceived as vowels. To disrupt formant constancy, the carrier frequency of each subsequent damped sinusoid was randomly selected from eight possible formant frequencies used to produce regular vowels, with randomization applied separately for each formant (frequency range: formant 1 = 270 - 1300 Hz; formant 2 = 850 - 2260 Hz; formant 3 = 1750 - 3000 Hz; formant 4 = 3300 - 5500 Hz). Both periodic and non-periodic sounds with disrupted formant structure were perceived as noises rather than vowels (i.e. non-vowels). The experiment included four stimulus types: (1) periodic vowels (/a/, /i/, /o/); (2) non-periodic vowels (/a/, /u/, /e); (3) three variants of periodic non-vowels and (4) three variants of non-periodic non-vowels. The spectral composition of these stimuli is presented in our previous paper (Orekhova et al., 2024).

Two hundred seventy stimuli from each of the four classes were presented, with three stimulus variants equally represented within each class (N = 90). All stimuli were presented in random order. Each stimulus lasted 812 ms, including 10 ms rise and fall time. The interstimulus intervals (ISIs) were randomly chosen from a range of 500 to 800 ms.

The non-periodic non-vowels were used as control stimuli. We examined how P100m characteristics change between test conditions (“periodic vowels”, “non-periodic vowels”, “periodic non-vowels”) and control condition (“non-periodic non-vowels”) and between TD and ASD groups.

### 2.3 Procedure

Children were instructed to watch a silent video (movie or cartoon) of their choice and ignore the auditory stimuli. Stimuli were delivered binaurally via plastic ear tubes inserted into the ear channels. The tubes were attached to the MEG helmet to prevent possible noise from their contact with the subject’s clothing. The stimulus mean intensity was set at 90 dB SPL. The experiment included three blocks of 360 trials each, with each block lasting approximately 9 minutes. Short breaks were introduced between the blocks. When necessary, a parent/caregiver remained with the child in the magnetically shielded room during the recording session.

### 2.4 MRI data acquisition and processing

In all participants with ASD and in 28 TD participants T1-weighted 3D-MPRAGE structural image was acquired on a Siemens Magnetom Verio 3T scanner (Siemens Medical Systems, Erlangen, Germany) using the following parameters: [TR 1780 ms, TE 2.78 ms, TI 900 ms, FA 9°, FOV 256L×L256Lmm, matrix 320L×L320, 0.8 mm isotropic voxels, 224 sagittal slices]. In 9 TD subjects MRIs were acquired at a 1.5T Philips Intera. In 2 TD subjects 1.5T GE Brivo MR355/MR360 was used. Cortical reconstructions and parcellations were generated using FreeSurfer v.7.4.1 (Dale et al., 1999; Fischl et al., 1999).

### 2.5 MEG data acquisition, preprocessing and source localization

MEG data were recorded using Elekta VectorView Neuromag 306-channel MEG detector array (Helsinki, Finland) with a 0.1 - 330 Hz bandpass filters and 1000 Hz sampling frequency.

Group differences in P100m parameters can be affected by differences in signal-to-noise ratio, which may be lower in autistic children. To control for this factor, we 1) monitored children’s head position during MEG recording and included in the analysis only those trials where head position and motion were below certain thresholds and 2) equalized the number of trials between groups (see below).

Bad channels were visually identified and labeled, after which the signal was preprocessed using MaxFilter software (v.2.2) to reduce external noise via the temporal signal-space separation method (tSSS) and to compensate for head movements by repositioning the head in each time point to an “optimal” common position. This position was determined individually for each participant as the position that resulted in the smallest average shift across all data epochs following motion correction.

Further preprocessing steps were performed using MNE-Python software (v.1.7.1) (Gramfort et al., 2013). The data were filtered using notch-filter at 50 and 100 Hz and a 110 Hz low-pass filter. Periods in which peak-to-peak signal amplitude exceeded thresholds of 7e-12 T for magnetometers or 7e-10 T/m for gradiometers, as well as “flat” segments where signal amplitude was below 1e-15 T for magnetometers or 1e-13 T/m for gradiometers were automatically excluded from further processing. To correct for cardiac and eye movement artifacts, we recorded ECG, vEOG, and hEOG and applied the signal-space projection (SSP) method. We then excluded from analysis data segments where head rotation exceeded a threshold of 10 degrees/s along any of the three space axes, head velocity exceeded 4 cm/s in 3D space, or head position deviated by more than 10 mm from the origin position in 3D space. The data were epoched from -0.2 s to 1 s relative to stimulus onset. The mean number of artifact-free data epochs was initially higher in the TD participants (TD: 997 vs ASD: 927, p < 0.05). To equalize these numbers, we randomly removed 70 epochs for each TD participant. The resulting mean number of clean epochs for each subject and stimulus type was 231 (range 141 - 340) for TD children and 231 (range 131 - 347) for children with ASD. The epoched data were averaged separately for each of the four experimental conditions and then baseline corrected by subtracting the mean amplitude in -200 - 0 ms prestimulus interval.

To obtain the source model, the cortical surfaces reconstructed with the Freesurfer were triangulated using dense meshes with about 130,000 vertices in each hemisphere. The cortical mesh was then resampled to a grid of 4098 vertices per hemisphere, corresponding to an average distance of about 4.9 mm between adjacent source points on the cortical surface.

To compute the forward solution, we used a single-layer boundary element model (inner skull). Source reconstruction of the averaged event-related fields was performed using the standardized low-resolution brain electromagnetic tomography (sLORETA) (Pascual-Marqui, 2002). Noise covariance was estimated from the time interval of -200 to 0 ms relative to stimulus onset.

### 2.6 Event related fields (ERF) analysis

#### 2.6.1 Sensor-level analysis of magnetic field distribution

To visualize magnetic field distribution on the heard surface, all subjects’ data were retransformed to a common standard head position (0, 0, 45 mm). RMS metric was computed based on the signal from all gradiometer sensors, separately for four types of stimuli (periodic vowels, periodic non-vowels, non-periodic vowels and non-periodic non-vowel). Grand average field distributions corresponding to the earliest RMS peak (P100m) and the later sustained negativity (SN) were plotted for TD and ASD groups (Figure 1).

**Figure 1.**
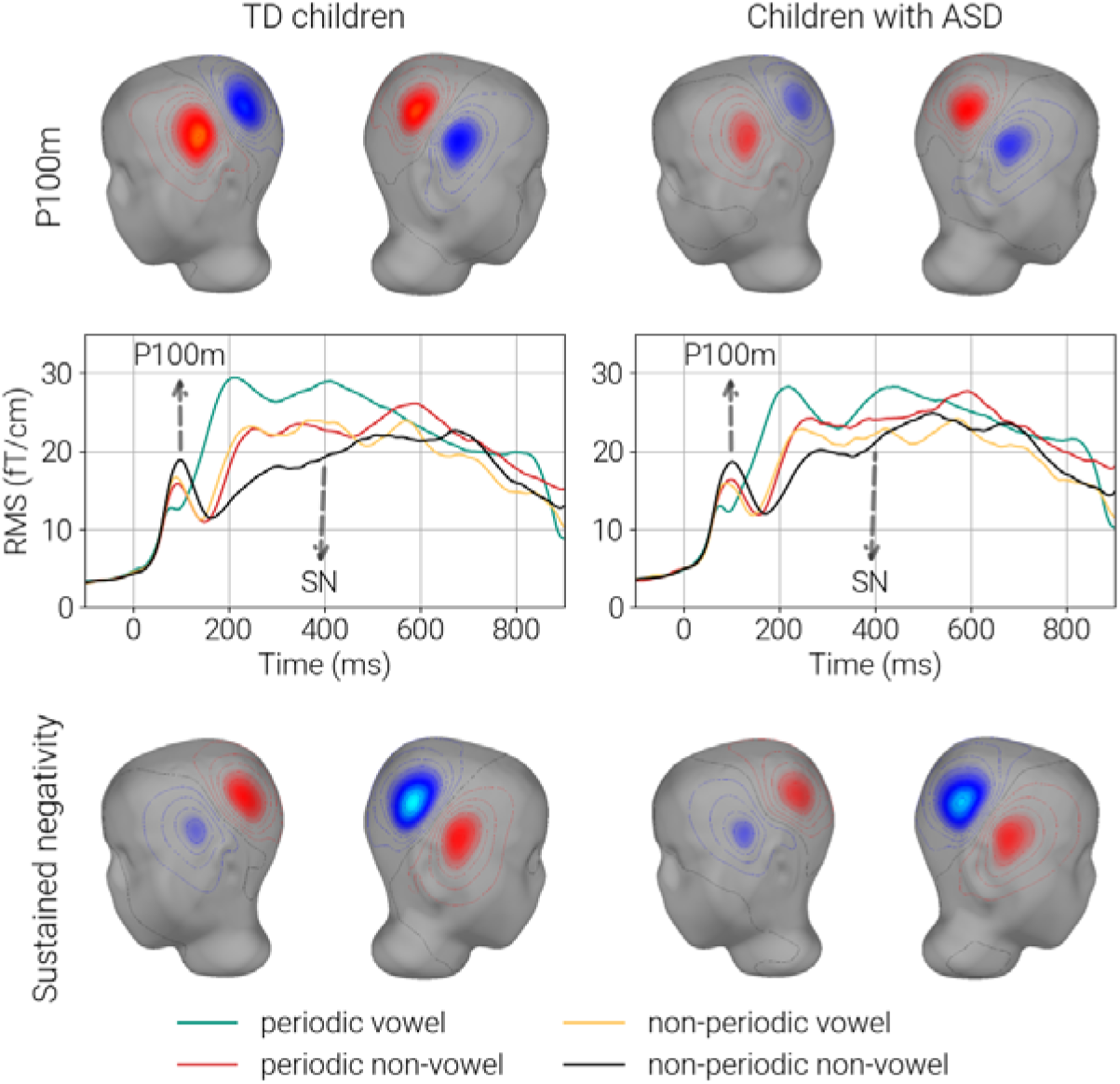
Magnetic Field Distribution. *Upper and lower panels*: Grand-averaged magnetic field maps derived from gradiometer data and projected onto the model head surface, shown separately for the TD and ASD groups. Maps display the control condition (non-periodic non-vowel), at 100 ms (P100m; upper panel), and 400 ms (SN; lower panel). *Middle panel*: Grand average root mean square (RMS) response waveforms. The horizontal axis zero point corresponds to stimulus onset (800-ms in duration). RMS values were computed across all gradiometer channels. Test conditions are represented by orange, green, and red lines; the control condition (non-periodic non-vowel) is shown in black.

#### 2.6.2 Detection of P100m component

The amplitude and latency of the P100m component was defined in the source space using each subject’s individual MRI–based brain model. The region of interest (ROI) was restricted to the superior surface of the temporal lobe, which includes functionally specialized auditory cortical areas A1, 52, LBelt, PBelt, MBelt, A4, TA2 (Glasser et al., 2016). To determine individual amplitudes and latencies of the P100m component, we undertook the two-steps procedure.

First, for each subject, we calculated *the grand averaged waveform*. To this end, vertex time courses within left and right ROIs were averaged first across four experimental conditions (periodic vowels; nonperiodic vowels; periodic non-vowels and non-periodic non-vowels) and then across all vertices within left and right ROIs using stc.extract_label_time_course. We used ‘mean_flip’ option, which ensures that the sign of the time series changes at vertices whose orientation differs from the dominant direction by more than 90°. The grand-average time course was subsequently low-pass filtered at 40 Hz using a fifth-order Butterworth filter. The grand-average P100m latency was approximated as the peak latency of the maximal positivity in the 50-140 ms time range.

Second, we calculated *single-vertex waveforms* by averaging the signal across all four conditions separately for each vertex in the ROI. We then computed correlations between single-vertex and grand-averaged waveforms in the P100m time window (50-140 ms) and identified vertices where the correlation exceeded 0.8 (Pearson’s R). From these vertices, we selected five sources with maximal amplitude within ±10 ms of the grand average P100 latency, which were considered to represent the subject’s P100m sources.

Inspection of individual time courses shown that this two-step procedure reduced the probability of detecting spurious sources and allowed characterization of the P100m component in all but one control participant. In this exceptional case, P100m was not detected in the right hemisphere.

Since the above P100m definition procedure assumed that P100m sources were identical across the four experimental conditions, we performed additional analysis to test this assumption. For this purpose, each subsects’ data were morphed into the FS-average template brain, the signal in the ROI was low-pass filtered at 40 Hz, and source orientations were adjusted according to the dominant direction. The MNI coordinates of the vertex with maximal amplitude of the signal in the P100m time window (50-140 ms) were identified separately for each of the four experimental conditions. Then, for each hemisphere and MNI coordinate (X, Y, Z), we performed mixed ANOVA with factors Condition and Group. No significant effects of Condition, Group or their interactions were found (all p > 0.1, uncorrected for multiple comparisons; see *Supplementary Table S1*). We therefore conclude that the assumption of common P100m sources across all conditions is valid.

#### 2.6.3 Estimation of sustained negativity (SN)

To estimate SN, the signal from individual P100m sources was averaged within 200-500 ms window, where SN was most prominent, separately for each condition.

#### 2.6.4 Statistical analysis

We applied Levene’s test to evaluate variance differences between TD and ASD groups. To examine the effect of Group (TD vs ASD), Condition (4 stimulus types) and Hemisphere on dependent variables (P100m latency and amplitude, SF amplitude), we conducted mixed ANOVA using type III sums of squares. In this approach, each fixed factor’s significance was tested after controlling for all other model factors. Partial eta-squared (*partial* η²) was computed to estimate effect sizes. Post hoc comparisons were conducted using the Wilcoxon signed-rank test, with false discovery rate (FDR) correction implemented via the Benjamini-Hochberg procedure.

If the variance of a parameter differed between the TD and ASD groups, we verified the ANOVA results using nonparametric tests.

For correlation analyses between component measures (e.g., latency or amplitude) and continuous variables (age, IQ), we used Spearman’s rank correlation. Partial or semi-partial Spearman correlation coefficients were computed using the ‘partial_corr’ function from the Python ‘pingouin’ package. When correlation directions were consistent across all conditions (e.g. P100m amplitudes–age correlations), we applied the Empirical Brown’s Method (EBM) to combine p-values (pcomb) (Poole et al., 2016) and calculated averaged correlation coefficient (rmean). Fisher’s z-transformation was employed for statistical comparison between correlation coefficients across groups or conditions.

## 3. Results

### 3.1 Sensor-level analysis of the auditory ERFs

Through visualizing magnetic field distribution on the head surface model, we observed that signals at 100 ms (P100m) and 400 ms (SN) displayed opposite polarities (Figure 1, lower and upper panels). Relative to control stimuli, stimuli characterized by temporal regularity (periodic non-vowels), formant structure (non-periodic vowels), or their combination (periodic vowels) evoked a transient RMS amplitude reduction around 100 ms (P100m time window), followed by a prolonged RMS increase persisting beyond 400 ms. The RMS amplitude reflects the average magnetic field strength independent of polarity, making it insensitive to the direction of state-dependent differences in cortical currents. Both the decreases and increases in RMS in response to vowel stimuli – compared to control non-periodic sounds lacking formant structure – reflect a sustained negative current shift in superior temporal cortex neural activity (Orekhova et al, 2024; Fadeev et al., 2024). To account for current polarity, we conducted further analysis in the source space.

### 3.2 Source-level analysis

#### 3.2.1 P100m localization

The P100m component was identified in both hemispheres of all participants except one TD child, in whom it was not detected in the right hemisphere. Figure 2A shows the localization of the vertices with maximal P100m amplitude in individual subjects, while Figure 2B presents the group medians. No group differences in the localization of the maximal P100m vertex were found for any MNI coordinates in either hemisphere (Mann-Whitney U test, all p > 0.069). In both groups, the median MNI coordinates corresponded to the primary auditory cortex (ASD: left hemisphere X=-49, Y=-23, Z=4, right hemisphere X=52, Y=-18, Z=3; TD: left hemisphere X=49, Y=-22, Z=2, right hemisphere X=52, Y=-19, Z=3) (Figure 2 B,C).

**Figure 2.**
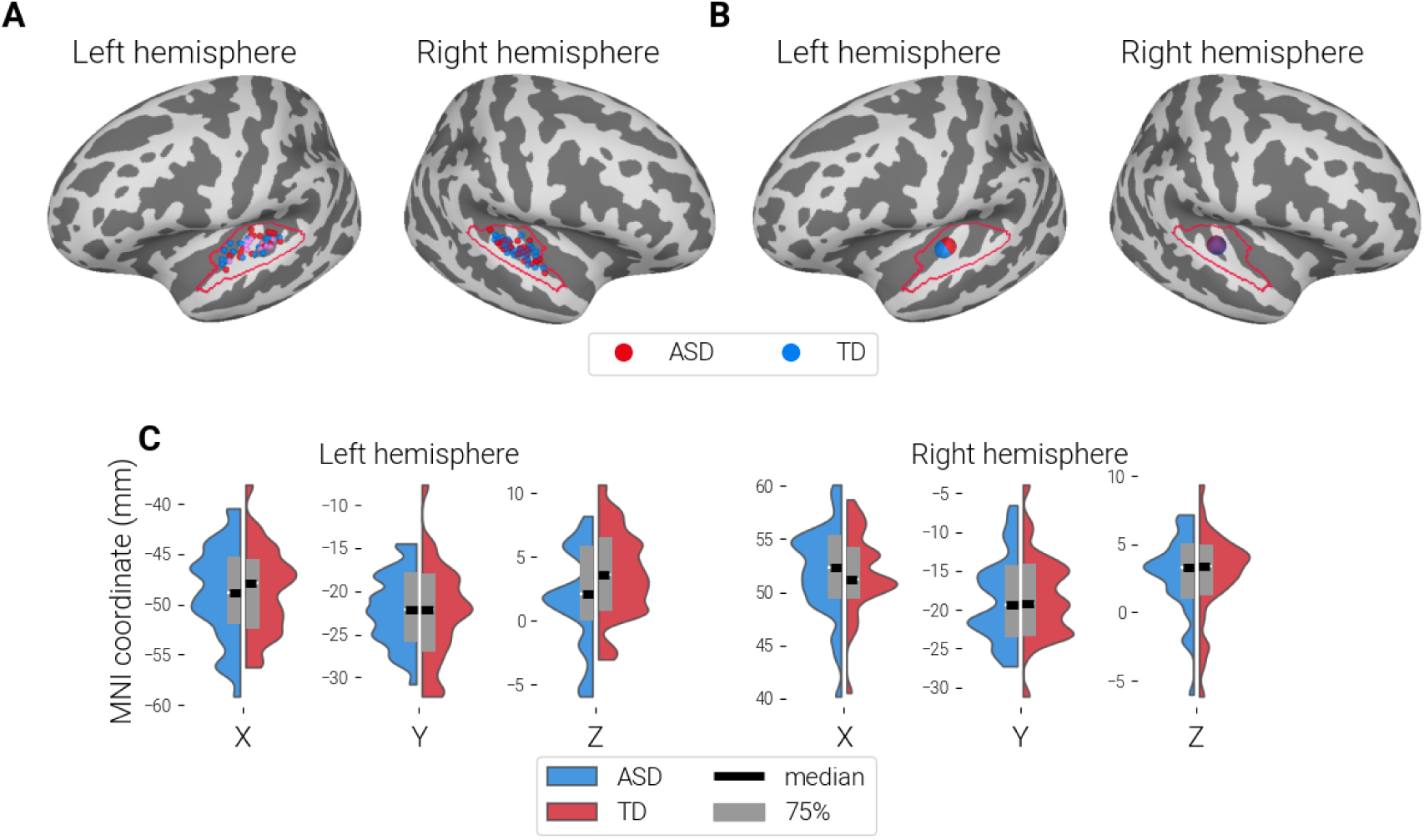
Localization of P100m Sources. **A:** Visualization of individual P100m ‘maximal vertices’ on the inflated surface of the FSaverage brain. Red dots represent children with ASD, while blue dots indicate TD children. The red lines delineate the regions of interest (ROIs) used to detect the P100m component. **B:** Group median coordinates for P100m localization. Red markers correspond to children with ASD, and blue markers represent TD children. **C:** Violin plots illustrating the MNI coordinates (X, Y, Z) for P100m localization in the left and right hemispheres for both TD and ASD groups. Each plot displays the median and the 75% interquartile range.

#### 3.2.2 P100m latencies and amplitudes in TD children and children with ASD

The mean P100m parameter values for TD and ASD children are presented in Table 2. No significant group differences were found in P100m latencies or amplitudes for any stimulus type in either hemisphere (Student T-test, all p > 0.25).

**Table 1.** Characteristics of the samples.

|  | Mean ± SD (range) |  | Student T-test | p |
| --- | --- | --- | --- | --- |
|  | *TD* | *ASD* |  |  |
| Age | Mean = 9.8 ± 1.8<br>(7.3 - 13.0)<br>N = 39 | Mean = 10.2 ± 1.7<br>(6.9 - 12.8)<br>N = 35 | T = 0.87 | 0.39 |
| MPI IQ<br>(KABC-II) | Mean = 116.2 ± 13.3<br>(85 - 138)<br>N = 39 | Mean = 83.1 ± 15.8<br>(54 - 128)<br>N = 34 | T = 9.7 | < 0.0001 |

N - number of participants for whom information is available.

**Table 2.** P100m amplitudes and latencies in TD and ASD groups. Mean ± standard deviation and range (in parenthesis) are given. Mann-Whitney U test was used for group comparisons.

|  | Periodic vowels |  | Non-periodic vowels |  | Periodic non-vowels |  | Non-periodic non-vowels |  |
| --- | --- | --- | --- | --- | --- | --- | --- | --- |
|  | LH | RH | LH | RH | LH | RH | LH | RH |
|  | Latency P100m |  |  |  |  |  |  |  |
| TD | 83±12,<br>(62-114) | 75±10,<br>(53-100) | 93±13,<br>(70-115) | 85±10,<br>(62-108) | 94±12,<br>(71-121) | 86±12,<br>(57-117) | 99±13,<br>(80-131) | 91±10,<br>(69-111) |
| ASD | 81±14,<br>(62-111) | 73±10,<br>(56-102) | 92±17,<br>(67-124) | 84±14,<br>(63-127) | 97±18,<br>(64-128) | 87±17,<br>(63-132) | 100±19,<br>(64-131) | 93±16,<br>(63-128) |
| M-W test | Z=-1.14,<br>p=0.25 | Z=-0.6,<br>p=0.55 | Z=-0.68,<br>p=0.5 | Z=-0.77,<br>p=0.44 | Z=-0.51,<br>p=0.61 | Z=-0.35,<br>p=0.73 | Z=-0.25,<br>p=0.8 | Z=-0.38,<br>p=0.71 |
|  | Amplitude P100m |  |  |  |  |  |  |  |
| TD | 9.69±4,<br>(3-21) | 7.56±3,<br>(1-14) | 12.6±5,<br>(5-28) | 10.3±4,<br>(1-19) | 12.0±5,<br>(5-26) | 9.72±4,<br>(2-18) | 13.6±6,<br>(5-30) | 11.48±5,<br>(2-21) |
| ASD | 8.84±4,<br>(1-17) | 8.1±4,<br>(2-16) | 10.91±4,<br>(3-20) | 10.37±5,<br>(2-24) | 10.83±4,<br>(2-20) | 10.47±5,<br>(2-22) | 12.77±5,<br>(2-30) | 11.91±5,<br>(1-24) |
| M-W test | Z=-0.65,<br>p=0.52 | Z=-0.59,<br>p=0.55 | Z=-0.91,<br>p=0.36 | Z=-0.09,<br>p=0.93 | Z=-0.28,<br>p=0.78 | Z=-0.51,<br>p=0.61 | Z=-0.06,<br>p=0.95 | Z=-0.43,<br>p=0.67 |

\*The data were available for all subjects ( $N_{td} = 39$ , $N_{asd} = 35$ ) in the left hemisphere and for all but one subject ( $N_{td} = 38$ , $N_{asd} = 35$ ) in the right hemisphere.
\*\* The p values are given without corrections for multiple comparisons

#### 3.2.3 Effect of age on latency and amplitude of P100m

Previous studies have shown that P100m latency decreases with age (Sharma et al., 2015), particularly in the right hemisphere (Roberts et al., 2019). To examine whether this developmental decrease in P100m latency was present in our study, we performed correlation analyses (Figure 3).

**Figure 3.**
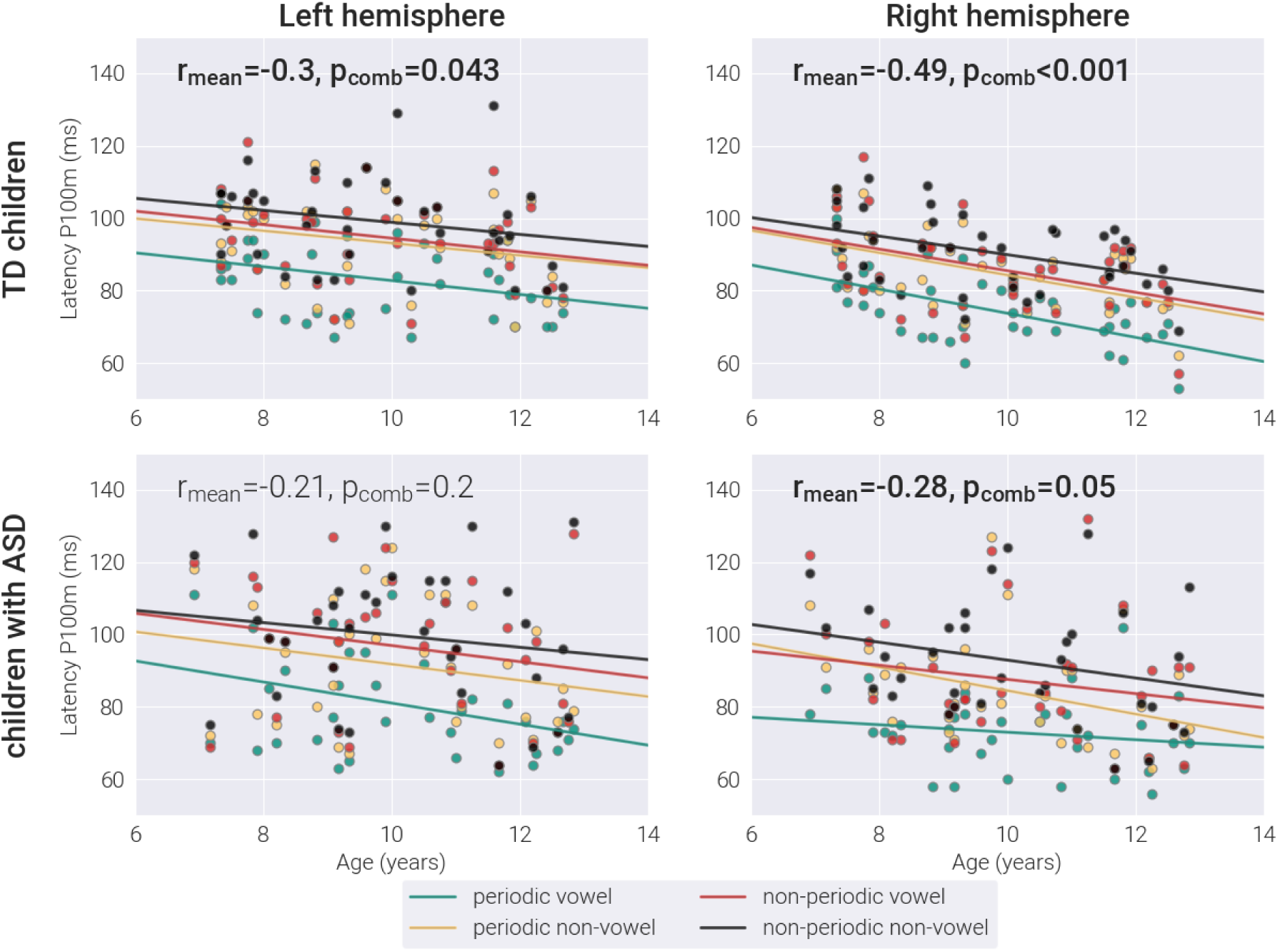
Correlation between P100m latency and age. Top panel: TD children; bottom panel: children with ASD. Colors denote different experimental conditions. Regression lines indicate an age trend for each stimulus type. Spearman r and the corresponding p-value were calculated using Empirical Brown’s Method.

P100m latency decreased significantly with age in TD children (LH: r_mean_ = -0.3, p_comb_ = 0.034, RH: r_mean_ = -0.49, p_comb_ < 0.0001). In children with ASD, this decrease was non-significant in the left hemisphere and marginally significant in the right hemisphere (LH: r_mean_ = -0.21, p_comb_ = 0.202, RH: r_mean_ = -0.28, p_comb_ = 0.05). However, no significant between-group differences were found in correlation coefficients (z-Fisher’s all p > 0.378).

Although correlations between P100m amplitude and age were predominantly negative in both groups (ranging from -0.06 to -0.18 across hemispheres and stimulation conditions in the combined sample), none reached statistical significance (all p> 0.13 for the combined sample).

#### 3.2.4 The time course of current in the P100m sources

Comparison of current time courses in P100m cortical sources between control and test stimuli revealed that periodicity, formant structure, or their combination produced significant current negativization started earlier than 100 ms, i.e. prior to the P100m peak (Figure 4). The enhanced negativity in response to test versus control condition – termed sustained processing negativity (SPN, Fadeev et al., 2024) - may mediate stimulus characteristic effect on P100m amplitude and latency. In subsequent sections, we examine how auditory regularities (periodicity and format composition) affect P100m latency and amplitude in children with ASD and TD participants.

**Figure 4.**
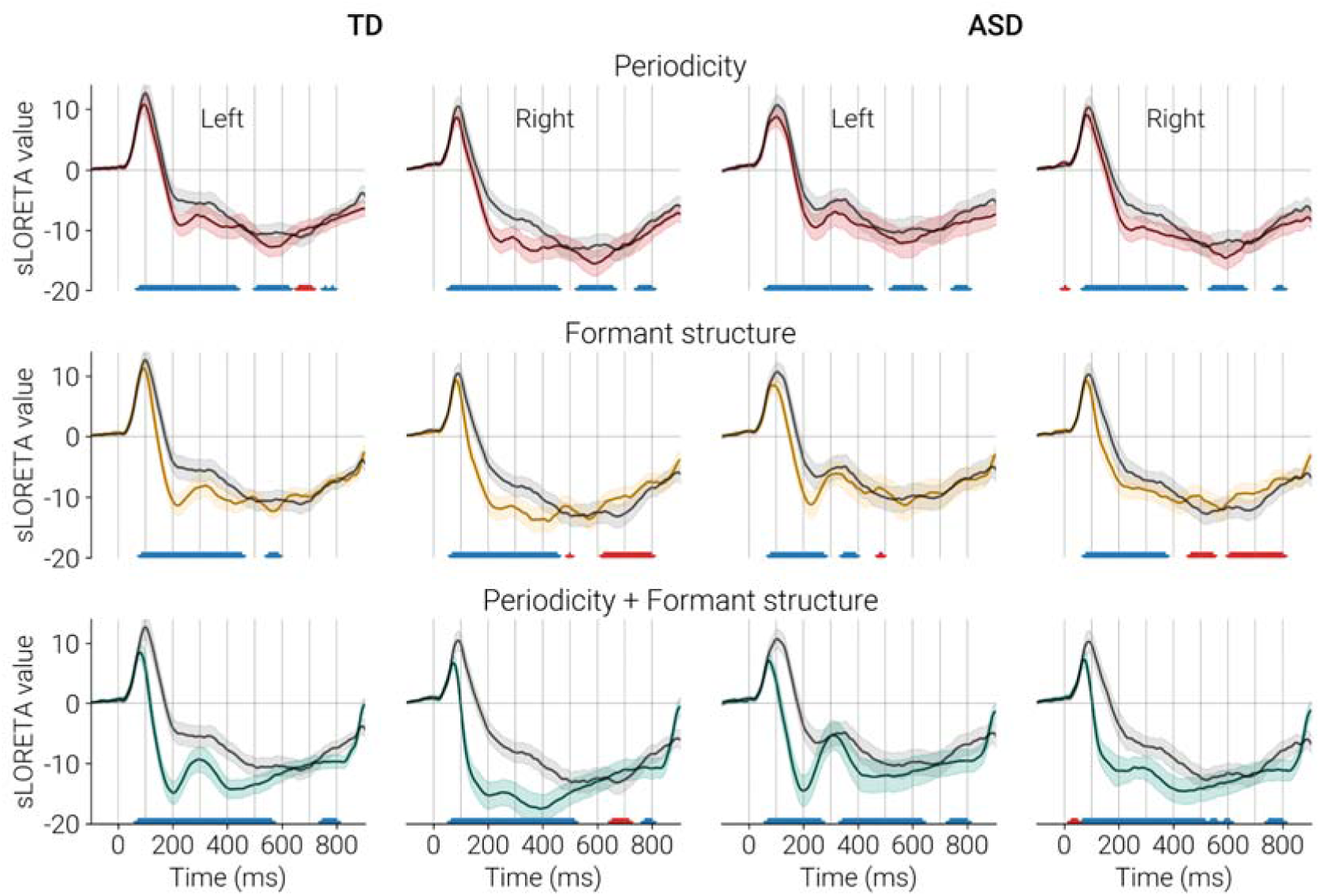
Grand-average time courses of auditory evoked responses in left and right P100m sources for TD children and children with and ASD. *Top Panel:* Periodicity effect (periodic non-vowel stimuli). *Middle Panel:* Formant structure effect (non-periodic vowel stimuli). *Bottom Panel:* Combined periodicity and formant structure effect (periodic vowel stimuli). Test conditions are shown as orange, green, and red lines; the control condition (non-periodic non-vowel) is indicated by the black line. Asterisks denote significant point-by-point differences between test and control conditions (paired *t*-test, *p* < 0.05, FDR-corrected). Blue: test<control, red test>control. Shaded areas represent 95% confidence intervals. Note the prolonged increase in negativity in response to test versus control that corresponds to sustained processing negativity (SPN).

#### 3.2.5 Influence of stimulus characteristics on P100m latency and amplitude

For P100m latency, Levene’s test indicated significantly greater variance in the ASD compared to the TD group in the left hemisphere for periodic non-vowels (F = 5.58, p = 0.021), non-periodic vowels (F = 5.74, p = 0.019) and non-periodic non-vowels (F = 4.76, p = 0.032) and in the right hemisphere for non-periodic non-vowels (F = 6.01, p = 0.016).

The mixed ANOVA revealed a significant main effect of Condition (F(3, 213) = 136.7, G-G epsilon = 0.67, p<0.0001, *partial* η² = 0.66), which was confirmed by significant Friedman ANOVA results (Left hemisphere: Chi Sqr. (N=74, df = 3) = 154,6, p<0.0001; Right hemisphere: Chi Sqr. (N=73, df = 3) = 151,2, p<0.0001). Post-hoc analysis using the Wilcoxon matched-pairs test showed that P100m latency was significantly longer in control versus test conditions bilaterally (all p’s < 0.0001, Figure 5; top panel).

**Figure 5.**
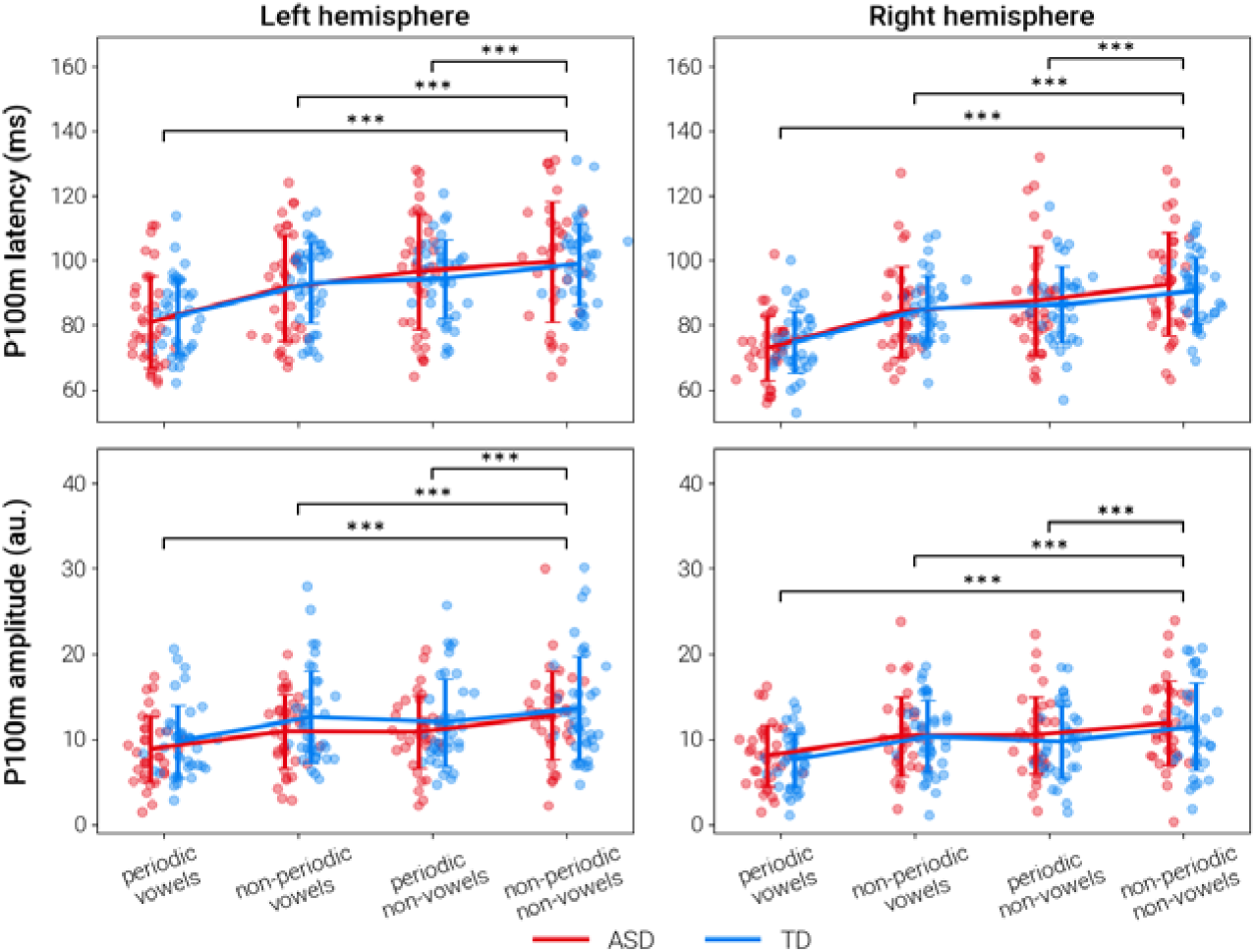
Individual P100m latency and amplitude values in TD and ASD groups. *Top panel:* P100m latency; *Bottom panel:* P100m amplitude. Error bars indicate standard deviations. *** p < 0.0001 (Wilcoxon signed-rank test, FDR-corrected for multiple comparisons between control versus test conditions).

A significant effect of Hemisphere was also observed (F(1, 71) = 31.7, p < 0.0001, ηp² = 0.31), with shorter P100m latencies in the right versus left hemisphere (Wilcoxon matched pairs test, for all conditions p<0.0001). No significant effect of Group or its interactions with Condition and Hemisphere were found for P100m latency (Table 3).

**Table 3.** Group, Condition and Hemisphere effects on P100m latency. Results of mrANOVA are presented with Greenhouse-Geisser correction.

| Fixed factor | numDF | denDF | F value | p value | *partial* $\eta^2$ |
| --- | --- | --- | --- | --- | --- |
| Group | 1 | 71 | 0.01 | 0.925 | 0.00 |
| **Condition** | **3** | **213** | **136.7** | **<0.0001** | **0.66** |
| **Hemisphere** | **1** | **71** | **31.7** | **<0.0001** | **0.31** |
| Group:Condition | 3 | 213 | 1.93 | 0.513 | 0.03 |
| Group:Hemisphere | 1 | 71 | 0.06 | 0.81 | 0.00 |
| Condition:Hemisphere | 3 | 213 | 0.18 | 0.90 | 0.00 |
| Group:Condition:Hemisphere | 3 | 213 | 0.16 | 0.93 | 0.00 |

No violation of the homogeneity of variance assumption was found for P100m amplitudes (Levene’s test, all p > 0.15). rmANOVA revealed a strong effect of Condition (F(3, 213) = 146.9, p < 0.0001, G-G epsilon = 0.81, *partial* η² = 0.67). Post-hoc analysis has shown that P100m amplitude was significantly higher in control than test conditions bilaterally (all p < 0.0001, Figure 5; bottom panel). A significant Hemisphere effect was also observed (F(1, 71) = 5.10, p = 0.027, *partial* η² = 0.07), with higher amplitudes in the left versus right hemisphere. No significant effects of the factor Group or its interactions with Condition and Hemisphere were found for P100m amplitude (Table 4).

**Table 4.** Group, Condition and Hemisphere effects on P100m amplitude. Results of mrANOVA are presented with Greenhouse-Geisser correction.

| Fixed factor | numDF | denDF | F value | p value | *partial* $\eta^2$ |
| --- | --- | --- | --- | --- | --- |
| Group | 1 | 71 | 0.17 | 0.68 | 0.00 |
| Condition | 3 | 213 | 146.9 | <0.0001 | 0.67 |
| Hemisphere | 1 | 71 | 5.10 | 0.027 | 0.07 |
| Group:Condition | 3 | 213 | 1.32 | 0.27 | 0.02 |
| Group:Hemisphere | 1 | 213 | 1.59 | 0.21 | 0.02 |
| Condition:Hemisphere | 3 | 213 | 0.11 | 0.95 | 0.00 |
| Group:Condition:Hemisphere | 3 | 213 | 0.32 | 0.981 | 0.00 |

In summary, both P100m latency and amplitude show a strong dependence on stimulus characteristics. Interhemispheric asymmetry is also evident, with the right hemisphere exhibiting shorter latencies and lower amplitudes compared to the left. These effects were consistent across both TD and ASD groups, with no significant between-group differences observed.

#### 3.2.6 Influence of stimulus characteristics on SN amplitude

Evidence from Figure 4, along with previous literature, suggests that the increase in SN associated with regular sound processing—also known as sustained processing negativity (SPN)—emerges in the auditory cortex prior to the peak of the P100m component (Hewson-Stoate et al., 2006; Molloy et al., 2019; Gutschalk & Uppenkamp, 2011; Fadeev et al., 2024). This enhanced SN may influence the transient P100m component of opposite polarity, even if their neural sources differ, due to the spatial field spread inherent in MEG/EEG recordings. Since our primary focus was the potential impact of SN on P100m, we estimated SN activity within the P100m sources by averaging the signal over the 200–500 ms window, thereby avoiding overlap with the preceding P100m response.

Our previous findings demonstrated SN enhancement by sound periodicity and/or formant composition. Here, we examined whether this pattern holds for SN estimated at P100m sources. rmANOVA revealed a significant Condition effect on SN amplitude (F(3, 213) = 97.1, G-G epsilon = 0.60, p < 0.0001, *partial* η² = 0.58) on the SN amplitude. Post-hoc analysis showed significantly less negative SN in control versus test conditions bilaterally (all *p* < 0.0001; Figure 6). A significant Hemisphere effect (F(1, 72) = 14.5, p = 0.0003, *partial* η² = 0.17) reflected greater SN negativity in the right versus left hemisphere (Table 5).

**Figure 6.**
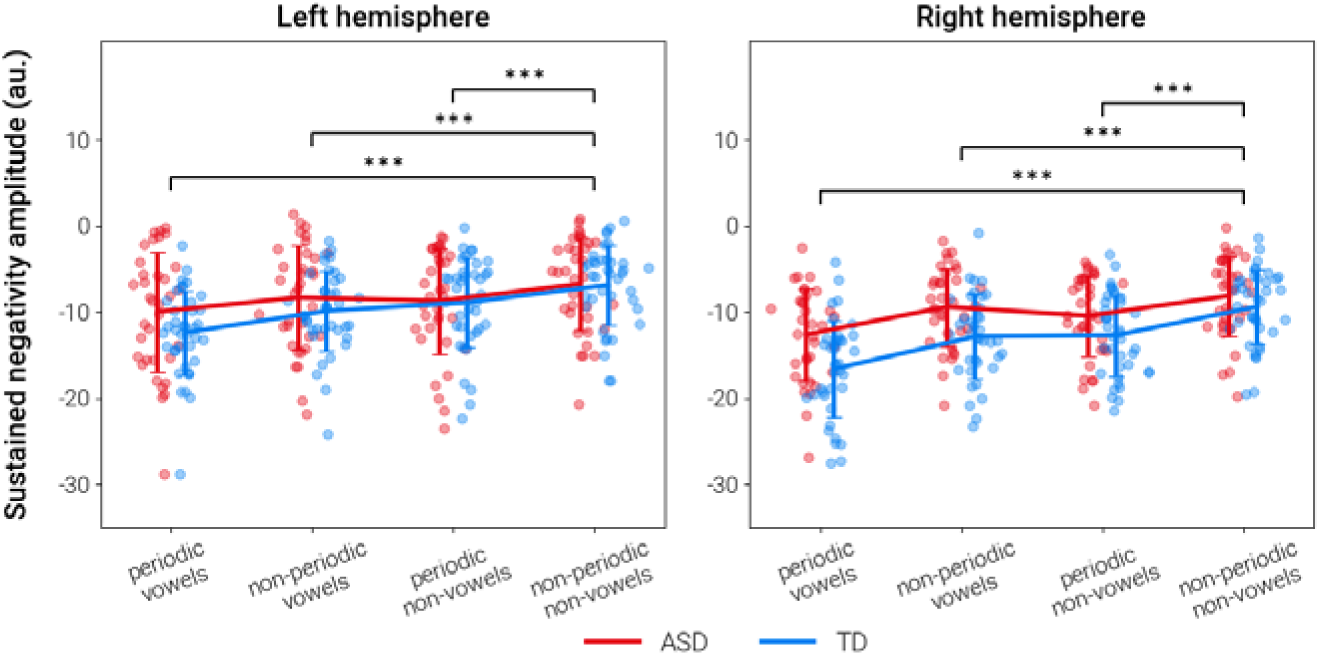
Individual SN amplitude values in TD and ASD groups. Error bars indicate standard deviations. *** p < 0.0001 (Wilcoxon signed-rank test, FDR-corrected for multiple control versus test condition comparisons).

**Figure 7.**
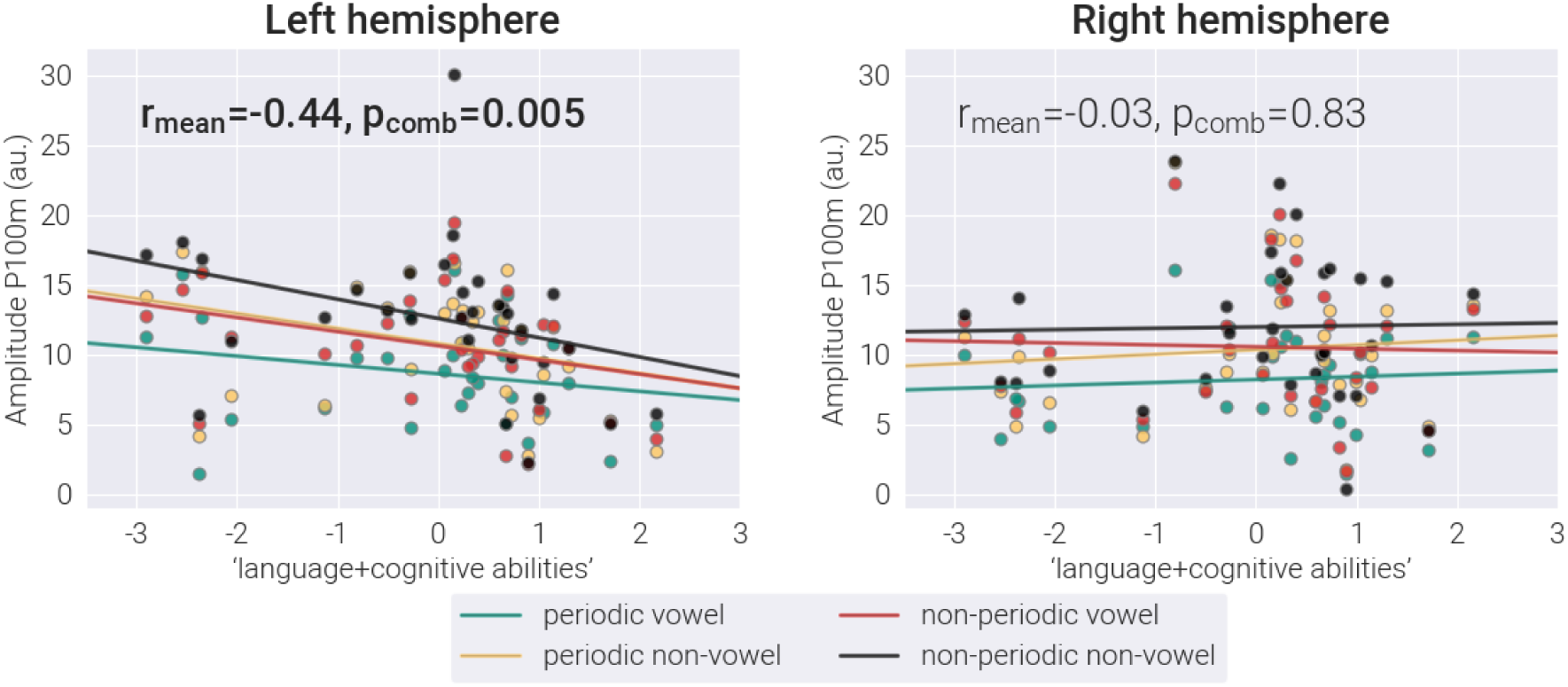
Age-corrected P100m amplitude associations with composite language and cognitive abilities in ASD children. Colored lines show condition-specific correlations with the PCA-derived score. The p_comb_ combined p-values estimated using Empirical Brown’s Method; r_mean_ - Spearman correlation coefficients, averaged across conditions.

**Table 5.** Group, Condition and Hemisphere effects on SN amplitude. Results of mrANOVA are presented with Greenhouse-Geisser correction.

| Fixed factor | numDF | denDF | F value | p value | *partial* $\eta^2$ |
| --- | --- | --- | --- | --- | --- |
| **Group** | **1** | **71** | **4.28** | **0.04** | **0.06** |
| **Condition** | **3** | **213** | **97.06** | **<0.0001** | **0.58** |
| **Hemisphere** | **1** | **71** | **14.53** | **<0.0001** | **0.17** |
| **Group\*Condition** | **3** | **213** | **7.06** | **<0.0001** | **0.09** |
| Group*Hemisphere | 1 | 71 | 1.51 | 0.22 | 0.02 |
| **Condition\*Hemisphere** | **3** | **213** | **5.67** | **0.001** | **0.07** |
| Group*Condition*Hemisphere | 3 | 213 | 0.45 | 0.71 | 0.01 |

Significant effects of Group and Group × Condition interaction were also observed (Table 5), reflecting reduced SN responses in the ASD group compared to TD participants for stimuli containing formant structure. The significant Hemisphere × Condition interaction indicated greater right-hemispheric dominance of SN for periodic versus non-periodic stimuli. These effects replicate those descrived in Fadeev et al. (2024) and Orekhova et al. (2024) using an alternative analytical approach and will not be discussed further here.

Overall, analysis of SN magnitude revealed condition-dependent differences that mirrored those observed for P100m magnitude. Compared to control conditions, test conditions consistently elicited reduced positivity (P100m) and enhanced negativity (SN) within their respective peak time windows. This effect was particularly pronounced for stimuli combining periodicity and formant structure. Notably, the condition eliciting the strongest P100m responses corresponded to the weakest SN magnitude, and vice versa, demonstrating an inverse relationship. A similar mirroring pattern was observed in hemispheric asymmetry: the right hemisphere exhibited stronger negative SN currents, while the left hemisphere showed greater positive P100m currents.

#### 3.2.7 P100m parameters, total language score and IQ in participants with ASD

Previous studies have reported associations between impaired language abilities and either delayed P100m responses (Yoshimura et al., 2013, 2021; Roberts et al., 2019) or increased P100m amplitudes (Roberts et al., 2019; Kautto et al., 2024). We examined whether general language abilities (assessed by the RuCLAB test) correlated with P100m parameters in ASD participants. Notably, the RuCLAB test is specifically designed to evaluate language abilities in children with speech and language impairments. Consequently, TD children frequently reach ceiling performance, precluding meaningful analysis of RuCLAB-P100m relationships in this group. For this reason, the analysis was restricted to children with ASD who had complete language and IQ scores available (N = 31).

Spearman partial correlation analysis (controlling for age) revealed associations between left hemisphere P100m amplitudes and total language scores in children with ASD. While individual stimulus conditions showed varying effect sizes (periodic vowel: r_part_ = -0.29, p = 0.114; non-periodic vowel: r_part_ = - 0.43, p = 0.017; periodic non-vowel: r_part_ = -0.32, p = 0.083; non-periodic non-vowel: r_part_ = -0.42, p = 0.022), the combined analysis revealed a significant negative correlation (r_mean_ = -0.37, p_comb_ = 0.027). This indicates that larger P100m amplitudes across stimulus conditions were associated with poorer language performance in children with ASD.

Semi-partial correlation analysis similarly revealed negative associations between age-corrected left P100m amplitudes and MPI IQ (periodic vowel: r_semipart_ = -0.19, p = 0.309; non-periodic vowel: r_semipart_ = - 0.33, p = 0.075; periodic non-vowel: r_semipart_ = -0.36, p = 0.049; non-periodic non-vowel: r_semipart_ = -0.45, p = 0.013; r_mean_ = -0.33, p_comb_ = 0.043).

Given the strong correlation between age-corrected language scores and MPI IQ in ASD children (r(31) = 0.50, p = 0.005), we performed principal component analysis (PCA) to extract their common factor. The first principal component (PC1), termed ‘composite language and cognitive ability’, explained 80% of variance and showed negative correlations with age-corrected P100m amplitudes across all stimulus conditions: (periodic vowel: r_semipart_ = -0.29 p = 0.115; non-periodic vowel: r_semipart_ = -0.48, p = 0.007; periodic non-vowel: r_semipart_ = -0.42, p = 0.022; non-periodic non-vowel: r_semipart_ = -0.57, p = 0.001; r_mean_ = - 0.44, p_comb_ = 0.005). Notably, the presence of temporal or spectral patterns in auditory stimuli did not significantly affect these correlations.

The same analysis conducted for right-hemisphere P100m amplitudes and for P100m latencies (in both hemispheres), revealed no significant effects (all p>0.22).

## 4. Discussion

We investigated how the P100m auditory response component in children was modulated by acoustic regularities, such as periodicity (pitch) and formant structure in synthetic vowels. Furthermore, we examined whether potential differences in P100m parameters between children with ASD and their TD peers were influenced by these regularities.

Our findings demonstrate that both temporal and spectral regularities reduce P100m amplitude and latency in both groups. At the group level, children with ASD showed no significant differences from TD children in P100m amplitude, latency, hemispheric lateralization, or stimulus-driven modulation patterns. However, the observed greater P100m latency variability in the ASD group suggests population heterogeneity and potential atypical auditory processing within the P100m time window for a subset of these children. Notably, while no group amplitude differences emerged, we found that stronger left-hemisphere P100m responses correlated with poorer cumulative language and cognitive performance measures in the ASD group.

In contrast to most previous developmental studies that employed dipole fitting analysis to localize the P100m (M50/P1m) component (e.g., Roberts et al., 2010, 2013, 2019, 2020; Edgar et al., 2015a, 2015b; Green et al., 2023; Matsuzaki et al., 2020; Yoshimura et al., 2013, 2016a, 2016b, 2021; Sano et al., 2024; Parviainen et al., 2019), we implemented a distributed source localization approach. Combining this method with individual MRI-based brain models, we successfully identified the P100m component in 100% of children in the left hemisphere and in 76 of 77 children in the right hemisphere. This detection rate meets or exceeds those reported in dipole source modeling studies (Roberts et al., 2010, 2020; Yoshimura et al., 2012, 2013, 2016a, 2016b, 2021; Sano et al., 2024). Consistent with previous developmental findings (Roberts et al., 2013; Sharma et al., 1997, 2015; Ponton, 2000), our data showed decreasing P100m latencies with age (Figure 3). Mirroring other reports (Edgar et al., 2015a; Yoshimura et al., 2021; Green et al., 2023), latencies were shorter in the right hemisphere compared to the left (Table 2), putatively reflecting faster maturation of the right-hemispheric auditory cortex (Parviainen et al., 2019). Similarly, we replicated the left-hemisphere amplitude dominance of P100m observed in prior pediatric studies (Yoshimura et al., 2013, 2021; Sano et al., 2024; Table 2). The median MNI coordinates localized the P100m source to Heschl’s gyrus in both groups, aligning with intracranial recording studies (Korzyukov et al., 2009; Liégeois-Chauvel et al., 1994, 2001; see Eggermont & Ponton, 2003 for review). The strong correspondence between our P100m characteristics and established literature confirms the reliability of our identification methodology.

The P100m and its EEG counterpart, the scalp-positive P100/P1, likely reflect contributions from three central nervous system pathways: the lemniscal pathway projecting to the primary auditory cortex (A1), the extralemniscal pathway terminating in secondary auditory areas that maintain dense reciprocal connections with A1, and reticular activating system (RAS) pathway. The selective preservation of P100m responses (in contrast to absent N1 components) observed in children with prolonged deafness suggests a strong contribution from non-specific thalamocortical projections, particularly those originating in the RAS, to the generation of this component (Eggermont & Moore, 2011).

Previous research has demonstrated that P100/P1 is modulated by preattentive arousal to novel auditory stimuli (Pratt et al., 2008) and its habituation (or gating) represents a neural correlate of the brain’s ability to filter out irrelevant, repetitive auditory input (Sinclair et al., 2017). In children, the enhanced P100/P100m amplitude at long interstimulus intervals suggests modulation by arousal responses to temporal novelty, whereas its attenuation during short-interval repetition reflects habituation mechanisms (Orekhova et al., 2013, Stroganova et al., 2013). While one might predict that perceptually salient periodic vowels (Fadeev et al., 2024) would elicit the strongest arousal and consequently the largest P100m responses, our findings revealed an opposite pattern: periodic vowels evoked the smallest P100m amplitudes, whereas unstructured control sounds produced the greatest responses. Intermediate P100m amplitudes were observed for sounds containing either temporal regularity alone (periodic non-vowels) or spectral regularity alone (non-periodic vowels) (Figure 5).

We propose that the observed inverse relationship between P100m amplitude and stimulus salience results from temporal overlap with SPN associated with the processing of temporal or frequency regularities. The SPN emerges during the P100m time window and persists for >400 ms post-stimulus onset (Figure 4; see also Orekhova et al., 2024; Fadeev et al., 2024). Notably, stimulus-related modulations of the positive-going P100m component show a directly opposite pattern compared to the negative-going SN measured in the 200–500 ms interval (Figures 5 and 6).

The progressive enhancement of SN - from control stimuli (lacking regularity) to test stimuli containing either temporal or spectral regularity, and ultimately to the ‘most regular’ test stimulus (periodic vowels) – likely reflects incremental activation of neural populations specialized for detecting auditory patterns characterizing ecologically relevant sounds such as vocalizations (Wang, 2018). Wang and colleagues (Bendor & Wang, 2005; Wang, 2007; Wang et al, 2008) described a class of auditory neurons responding to periodic sounds with sustained firing that is not timed to the onset of each individual sound pulse in a periodic sequence (so-called “unsynchronized neurons”). Similarly, neurons that respond tonically to certain frequency compositions characteristic of conspecific vocalizations (Tian et al., 2001) may account for the increase in SN in response to vowels devoid of periodicity. Hence, the response to periodic vowels may reflect the joint activity of these neuronal populations. Studies in humans (Malloy et al., 2019) and animals (Walker et al., 2011) show that auditory patterns are detected very rapidly by the brain and can begin modulating evoked responses as early as ∼40 ms after stimulus onset—well before the P100m reaches its peak.

Taken together, our findings suggest that the auditory cortex evoked activity during the P100m time window is mediated by at least two distinct neural processes: (1) a phasic surface-positive response that matures early, peaks around 100 ms post-stimulus in response to any detectable auditory input, and originates primarily from cortical layers IIIc and IV (Eggermont & Ponton, 2003; Eggermont & Moore, 2011; present findings); and (2) sustained surface-negative activity originating in supragranular layers II and III, which becomes more pronounced during the encoding of behaviorally relevant sounds that contain temporal regularities or distinct spectral characteristics (Wang, 2007). These processes exert opposing influences on P100m amplitude, and variability in either of them may contribute to previously reported group differences in P100m parameters between children with ASD and typically developing peers during complex speech processing (Yoshimura et al., 2013, 2016b; Sano et al., 2024).

Our study revealed no significant group differences in mean P100m amplitudes or latencies between children with ASD and TD children across all stimulus conditions. The absence of P100m amplitude differences in ASD is consistent with recent meta-analytic findings (Williams et al., 2020). However, the lack of latency differences contrasts with several previous studies that reported delayed P100m responses in ASD (Roberts et al., 2013, 2019, 2020; Matsuzaki et al., 2020; see also Williams et al., 2020). Several factors may account for this discrepancy. First, the pronounced heterogeneity within the ASD population is evident in our data: P100m latency variance was significantly greater in the ASD group compared to TD controls. Notably, for the control stimulus—which elicited the strongest P100m responses—the ASD group included both the shortest and longest latency values (Figure 5, upper panel). Second, stimulus characteristics may play a role. While previous studies reporting delayed P100m latencies in ASD typically used pure tones, we employed spectrally complex stimuli. Notably, Yoshimura and colleagues, who also used complex auditory stimuli (e.g., the Japanese syllable /ne/), reported either typical (Yoshimura et al., 2013) or even shorter (Yoshimura et al., 2016b, 2021) P100m latencies in ASD participants compared to TD controls.

Building on prior research demonstrating associations between P100m characteristics and language abilities (Yoshimura et al., 2013, 2021; Roberts et al., 2019; Kautto, 2024), we expected to observe similar relationships in our study. We focused this analysis exclusively on the ASD group, as TD children consistently achieved near-ceiling performance on the language assessment. Given the strong correlation between language performance and cognitive abilities in the ASD group, coupled with our limited sample size, we were unable to reliably dissociate these factors. Consequently, we employed a composite measure combining both language and cognitive abilities.

We found a significant inverse relationship between P100m amplitude and combined language-cognitive scores in autistic participants. This finding is consistent with Roberts et al. (2019), who reported increased P100m amplitudes in response to pure tones in minimally verbal children with ASD compared to their verbal peers. In our study, this association was most pronounced for left-hemisphere responses to non-periodic, non-vowel stimuli while failing to reach statistical significance for periodic vowels. The robust P100m responses evoked by non-periodic, non-vowel stimuli, likely reflecting an improved signal-to-noise ratio, may have enhanced the detection of the link between language-cognitive abilities and P100m amplitude. These results suggest that the observed association is not specific to the processing of acoustic characteristics of vowel sounds, but rather represents more general neural mechanisms that modulate P100m responses to different auditory stimuli.

The observed association between increased P100m amplitudes and lower language-cognitive abilities in ASD may have at least two, not mutually exclusive explanations.

First, those children with ASD who are more severely affected may exhibit delayed maturation of the transient N100 component, which originates from distinct auditory cortical circuitry that matures later than the P100-generating networks (Eggermont & Moore, 2011). Since the N100 typically emerges in a time window partially overlapping with P100 (Orekhova et al., 2013), its progressive maturation may attenuate the late P100m portion, ultimately shortening P100m latency with age. Although our paradigm, using short ISIs, did not elicit a distinct N100m peak (Budd et al., 1998; Čeponien et al., 1998), the underlying maturational processes may still influence P100m characteristics.

Second, more severely affected children with ASD may show deficient habituation/sensory gating, which could manifest as elevated P100m amplitudes. This is consistent with findings of P50/P100m habituation deficits in ASD (Ruiz-Martínez et al., 2020; Orekhova et al., 2008), where the severity of impairment correlates with the degree of intellectual disability (Orekhova et al., 2008).

Despite its negative association with the psychometric scores, the P100m amplitudes in children with ASD were within the typical range (Table 2). This lack of group difference may be related to the influence of additional factors associated with ASD, such as increased neural noise, which could act in the opposite direction by enhancing response variability and effectively reducing P100m amplitude (Milne, 2011; David et al., 2016; Raul et al., 2024).

Our findings carry important methodological implications for P100/P100m research in clinical populations. When studying the transient P100/P100m component, irregular auditory stimuli (e.g., noise bursts) appear superior for reliable component identification, as they evoke robust responses with favourable signal-to-noise characteristics. Conversely, patterned stimuli like vowels typically elicit attenuated P100/P100m amplitudes with poorer signal quality, potentially compromising accurate parameter estimation.

## 5. Conclusion

In summary, our findings reveal that P100m responses to temporally or spectrally regular auditory stimuli integrate at least two distinct neural mechanisms: (1) a phasic component reflecting sound detection and preattentive arousal, and (2) a sustained component involved in auditory pattern processing. Both processes may contribute to the variability in P100m characteristics observed across clinical populations.

While we found no significant group-level differences in P100m amplitudes or latencies between children with ASD and their TD peers, this does imply typical P100m generation in ASD. Notably, the ASD group exhibited substantially greater P100m latency variability than TD controls, suggesting neurobiological heterogeneity that may encompass distinct subgroups - some showing atypically prolonged latencies (consistent with Roberts et al., 2013, 2019; Matsuzaki et al., 2020) and others demonstrating atypically short latencies (consistent with Yoshimura et al., 2016b, 2021). Furthermore, in children with ASD, higher left-hemisphere P100m amplitudes to the auditory stimuli – particularly non-patterned sounds - were negatively correlated with language and cognitive performance. This association may indicate an adverse impact of atypical pre-attentive auditory arousal/habituation or delayed auditory cortex maturation on language and general cognitive functioning in this population.

These results underscore the complex relationship between auditory P100m responses and language impairments in ASD, which appears to emerge from the interplay of multiple neural mechanisms. Further studies that specifically target each underlying process and include diverse clinical populations are needed to determine whether this component of the auditory response can serve as a useful biomarker for ASD.

## Ethics Statement

The Ethics Committee of the Moscow State University of Psychology and Education approved this study. All children provided verbal assent to participate in the research, and their caregivers provided written informed consent for participation.

## Conflicts of Interest

Not applicable.

## Data Availability Statement

The de-identified individual-level raw data that supports this research, study materials, and analysis code used to generate the results are publicly available at https://openneuro.org/datasets/ds005234.

## Notes

https://openneuro.org/datasets/ds005234

### Competing Interest Statement

The authors have declared no competing interest.

### Summary of Updates

Corrected an error in the co-author name from Ilakai V Romero Reyes to Ilacai V Romero Reyes.

