## Supplementary Table S1 for "Vowel-like patterns modulate auditory P100m response but not its association with language abilities in children with ASD"

### Supplementary materials

Table S1 - Results of rmANOVA for MNI coordinates with factors Condition and Group. Values are presented with Greenhouse-Geisser correction.

|  | Left hemisphere, X axis | | | | | | |
| --- | --- | --- | --- | --- | --- | --- | --- |
|  | numDF | denDF | SSn | SSd | F value | p value | general η² |
| Group | 1 | 72 | 27.968 | 5110.564 | 0.39 | 0.532 | 0.004 |
| Condition | 2.63 | 189.03 | 35.556 | 2230.885 | 1.15 | 0.328 | 0.005 |
| Group:Condition | 2.63 | 189.03 | 20.033 | 2230.885 | 0.65 | 0.566 | 0.003 |
|  | Left hemisphere, Y axis | | | | | | |
|  | numDF | denDF | SSn | SSd | F value | p value | general η² |
| Group | 1 | 72 | 20.04 | 6664.049 | 0.22 | 0.643 | 0.002 |
| Condition | 3 | 216 | 8.323 | 3660.815 | 0.16 | 0.921 | <0.001 |
| Group:Condition | 3 | 216 | 14.254 | 3660.815 | 0.28 | 0.84 | 0.001 |
|  | Left hemisphere, Z axis | | | | | | |
|  | numDF | denDF | SSn | SSd | F value | p value | general η² |
| Group | 1 | 72 | 67.478 | 3136.337 | 1.55 | 0.217 | 0.014 |
| Condition | 2.61 | 187.85 | 51.618 | 1789.034 | 2.08 | 0.113 | 0.01 |
| Group:Condition | 2.61 | 187.85 | 8.327 | 1789.034 | 0.34 | 0.772 | 0.002 |
|  | Right hemisphere, X axis | | | | | | |
|  | numDF | denDF | SSn | SSd | F value | p value | general η² |
| Group | 1 | 72 | 0.444 | 5326.631 | 0.01 | 0.938 | <0.001 |
| Condition | 2.42 | 174.3 | 45.789 | 2597.375 | 1.27 | 0.286 | 0.006 |
| Group:Condition | 2.42 | 174.3 | 14.489 | 2597.375 | 0.4 | 0.709 | 0.002 |
|  | Right hemisphere, Y axis | | | | | | |
|  | numDF | denDF | SSn | SSd | F value | p value | general η² |
| Group | 1 | 72 | 5.572 | 8329.129 | 0.05 | 0.827 | <0.001 |
| Condition | 2.42 | 173.94 | 36.817 | 4669.253 | 0.57 | 0.6 | 0.003 |
| Group:Condition | 2.42 | 173.94 | 56.881 | 4669.253 | 0.88 | 0.435 | 0.004 |
|  | Right hemisphere, Z axis | | | | | | |
|  | numDF | denDF | SSn | SSd | F value | p value | general η² |
| Group | 1 | 72 | 116.482 | 2774.29 | 3.02 | 0.086 | 0.027 |
| Condition | 2.3 | 165.32 | 30.378 | 1501.412 | 1.46 | 0.234 | 0.007 |
| Group:Condition | 2.3 | 165.32 | 21.488 | 1501.412 | 1.03 | 0.367 | 0.005 |
